# Development and validation of a subject-specific integrated finite element musculoskeletal model of human trunk with ergonomic and clinical applications

**DOI:** 10.1101/2024.01.06.574467

**Authors:** Farshid Ghezelbash, Amir Hossein Eskandari, Aboulfazl Shirazi-Adl, Christian Larivière

## Abstract

**Background and Objectives:** Biomechanical modeling of the human trunk is crucial for understanding spinal mechanics and its role in ergonomics and clinical interventions. Traditional models have been limited by only considering the passive structures of the spine in finite element (FE) models or incorporating active muscular components in multi-body musculoskeletal (MS) models with an oversimplified spine. This study aimed to develop and validate a subject-specific coupled FE-MS model of the trunk that integrates detailed representation of both the passive and active components for biomechanical simulations.

**Methods:** We constructed a parametric FE model of the trunk, incorporating a realistic muscle architecture, personalized through imaging datasets and statistical shape models. To validate the model, we compared tissue-level responses with in vitro experiments, and muscle activities and intradiscal pressures versus in vivo measurements during various physical activities. We further demonstrated the versatility of the proposed personalized integrated framework through additional applications in ergonomics (i.e., wearing an exoskeleton) and surgical interventions (e.g., nucleotomy and spinal fusion).

**Results:** The model demonstrated satisfactory agreement with experimental data, showcasing its validity to predict tissue- and disc-level responses accurately, as well as muscle activity and intradiscal pressures. When simulating ergonomics scenarios, the exoskeleton-wearing condition resulted in lower intradiscal pressures (1.9 MPa vs. 2.2 MPa at L4-L5) and peak von Mises stresses in the annulus fibrosus (2.2 MPa vs. 2.9 MPa) during forward flexion. In the context of surgical interventions, spinal fusion at L4-L5 led to increased intradiscal pressure in the adjacent upper disc (1.72 MPa vs. 1.58 MPa), whereas nucleotomy minimally influenced intact disc pressures but significantly altered facet joint loads and annulus fibrosus radial strains.

**Conclusions:** The integrated FE-MS model of the trunk represents a significant advancement in biomechanical simulations, providing insights into the intricate interplay between active and passive spinal components. Its predictive capability extends beyond that of conventional models, enabling detailed risk analysis and the simulation of varied surgical outcomes. This comprehensive tool has potential implications for the design of ergonomic interventions and the optimization of surgical techniques to minimize detrimental effects on spinal mechanics.

## 1 Introduction

More accurate knowledge of the human spine is a critical prerequisite in designing more effective injury prevention programs, evaluation, rehabilitation, and treatment planning. Although experimental measurements (e.g., electromyography, intradiscal pressure, kinematics) provide valuable insights into spine biomechanics, they remain difficult to perform, costly, invasive and/or limited in predictive power (Beaudette et al., 2014; Ghezelbash et al., 2020b; Mannen et al., 2018, 2015; Nachemson and Morris, 1964; Reed et al., 2022). On the other hand, biomechanical models, once constructed, driven and validated by experimental data, are promising in performing comprehensive analyses to gain deeper understanding of human biomechanics, carrying out injury risk assessment, and designing novel therapeutic interventions.

Biomechanical models of the human trunk can be categorized into two general groups: 1-finite element (FE) passive spine models (Dreischarf et al., 2014; Stott and Driscoll, 2023) and 2- multi-body dynamics musculoskeletal (MS) models (Liu et al., 2023). Detailed FE models of a ligamentous spine (devoid of muscles) can accurately represent the structure and mechanics of the passive spine while overlooking muscles as active components (Schmidt et al., 2007a; Shirazi-Adl, 2006; Xu et al., 2017). On the other hand, MS models incorporate muscles along with a passive model and estimate their forces using optimization and/or electromyography (EMG) approaches. These latter models with the exception of few recent ones (Ebrahimkhani et al., 2022; Rajaee et al., 2021), have used simplistic representations (e.g., spherical joints with/without rotational springs, beams) of the passive spine (Eskandari et al., 2023a; Ghezelbash et al., 2015, 2018; Meng et al., 2015). Despite the crucial mechanical interaction between active (i.e., muscles) and passive (i.e., ligamentous spine) components, biomechanical models often make assumptions for the sake of computational efficiency when neglecting muscles in FE passive spine models or using simplified passive elements to simulate spinal joints in MS models. Such idealizations could adversely influence the accuracy of estimations (Ghezelbash et al., 2018, 2015; Meng et al., 2015). Furthermore, biomechanical models regularly use a single generic model with a unique input geometry, body weight and kinematics thus neglecting differences among individual subjects that also potentially affect results (Firouzabadi et al., 2021; Ghezelbash et al., 2016b, 2017; Ignasiak et al., 2018). Alternatively, combining a detailed coupled FE-MS model of the spine in the context of a subject-specific framework promises to resolve many fundamental limitations of earlier models, and to lay the groundwork for future developments.

Here, we aim to develop and validate a novel subject-specific MS model of the trunk that directly integrates a detailed FE passive model. A parametric model of a human trunk is initially constructed with the model parameters being adjusted based on available imaging datasets and statistical shape models in accordance with varying subject’s height, weight, age and sex. To validate the model, we compare computed tissue-level responses (under compression and pure moment) with *in vitro* experiments, and muscle activities and intradiscal pressures versus *in vivo* measurements during various physical activities. We further demonstrate the potential of the proposed personalized integrated framework through additional applications in ergonomics (i.e., wearing an exoskeleton) and surgical interventions (i.e., nucleotomy and spinal fusion).

## 2 Methods

### 2.1 Subject-specific FE-MS model

#### Passive spine FE model

A previously validated detailed FE model of the lumbar spine (T12 to S1) was used as the reference model (Rajaee et al., 2021; Shirazi-Adl, 1994a, 1994b). A novel scaling algorithm was introduced to individualize this FE model of the spine using regression equations and statistical models. In this reference passive spine model, collagen fibers (in the disc annulus fibrosus) from two adjacent layers were homogenized into a single layer forming a crisscross pattern with ±30° angles to the transverse plane. The collagen volume content and mechanical properties were adjusted along the annulus radial direction to mimic the reported changes in content and transition from the collagen type I (at outermost layers) to collagen type II (at innermost layers) (Ghezelbash et al., 2021; Shirazi-Adl et al., 1986). To reduce the computational cost, the disc nuclei were simulated as incompressible inviscid fluid-filled cavities (i.e., no shear resistance), and all bony vertebrae were simulated as a collection of two rigid bodies where T12-L5 vertebrae were divided into distinct anterior and posterior rigid bodies attached by deformable beams representing their pedicles (Shirazi-Adl, 1994a, 1994b). All materials were assumed to be elastic. For more details on material properties, see Supplementary Materials (Ghezelbash et al., 2020a).

To individualize the model, we parametrized the reference model (Figure 1) based on existing imaging studies and statistical shape models in accordance with subject’s sex, age, height and weight (Tang et al., 2022, 2019). Disc areas (anterior-posterior and medio-lateral disc diameters) were adjusted based on a subject’s sex and height (Tang et al., 2019), and segmental lordosis was scaled in accordance with a statistical shape model of the lumbar spine (Tang et al., 2022), Figure 1. Disc heights were assumed to be linearly proportional to the subject’s body height (Ghezelbash et al., 2016a). Nodal coordinates of posterior elements (e.g., facet articular surfaces, transverse processes, spinous processes) were modified proportional to the disc geometry (Figure 1). Collagen fiber network parameters (i.e., fiber area and angle) were adjusted to maintain the same volume fraction content and orientation within the respective intervertebral disc.

**Figure 1:**
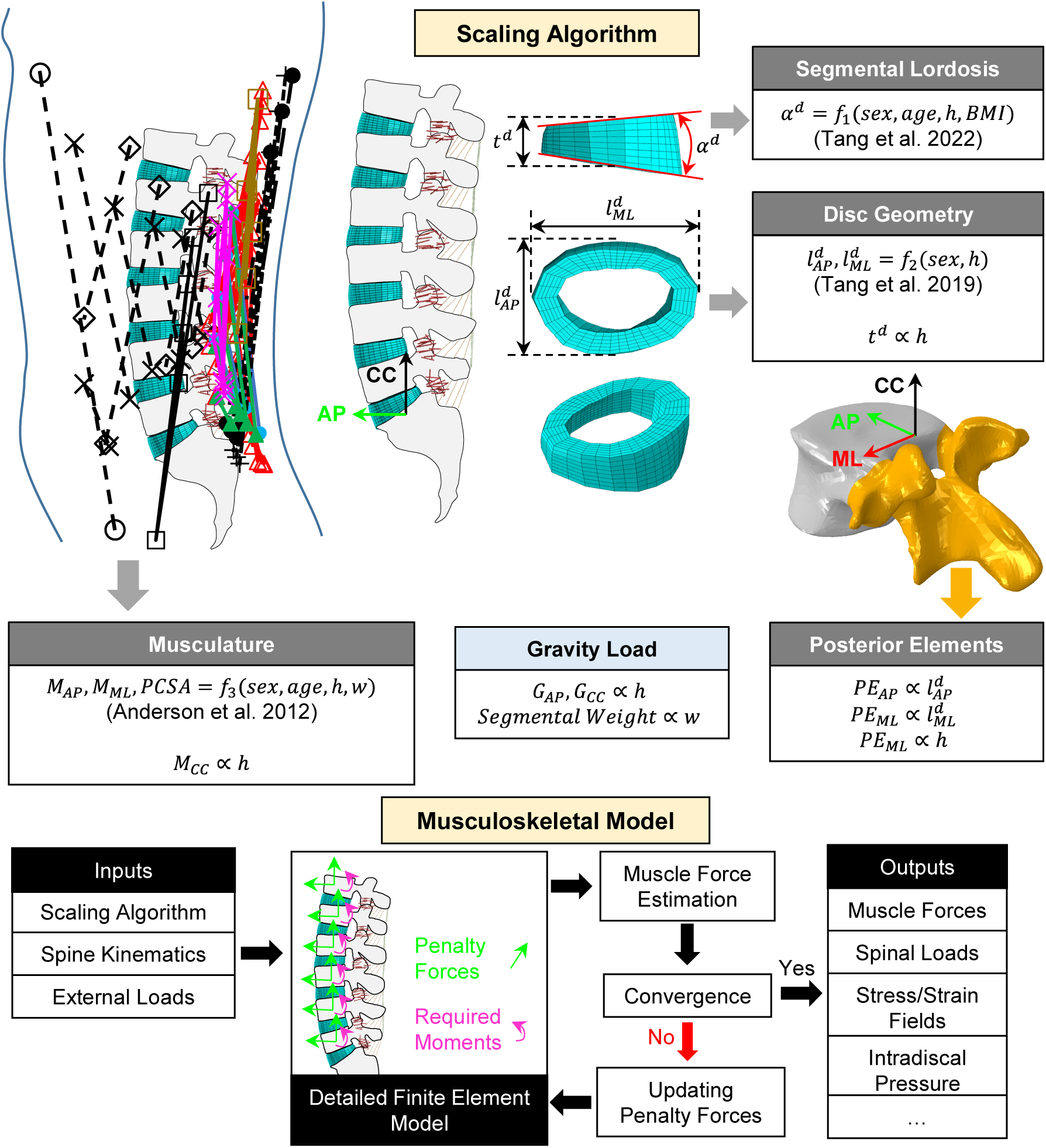
Schematics of the scaling algorithm and the subject-specific FE-MS model (*α*^d^: segmental lordosis; *t*^d^: mean disc height; *l*^d^: disc diameters; *PE*: nodal coordinates of posterior elements; *M*: muscle moment arm; *PCSA*: physiological cross sectional area; *G*: gravity load location; *h*: body height; *w*: body weight; *BMI*: body mass index; *AP*: anterior-posterior; *ML*: medio-lateral; *CC*: cranial-caudal).

#### Musculoskeletal model

The foregoing individualized FE model of the passive spine was integrated with a comprehensive musculature incorporating 126 sagittally-symmetric muscles (Eskandari et al., 2023b; Ghezelbash et al., 2016b). Muscle moment arms and cross-sectional areas were individualized with subject’s anthropometric parameters (i.e., sex, age, height and weight) (Anderson et al., 2012a); Figure 1. Segmental masses along the trunk height were proportionally scaled based on the subject’s body weight while their lever-arms were adjusted based on the subject’s body height (De Leva, 1996; Pearsall et al., 1996). In each task and after the individualization of the passive spine model, musculature and segmental masses; measured kinematics (i.e., segmental rotations at T12 to S1 levels) and external loads were applied into the model, and muscle forces were estimated (using the minimization of the sum of squared muscle stresses constrained by equilibrium equations at all levels-directions). Muscle forces were subsequently updated and added to the existing external loads in the next iteration. The analysis was repeated until convergence was reached (<5% changes in muscle forces) (Seth and Pandy, 2007).

### 2.2 Validation

The validation of the proposed personalized FE-MS model was carried out at different levels:

#### Annulus fibrosus tissue

We compared stress-strain responses of the annulus fibrous at the tissue level against experimental studies under uniaxial (along fiber directions) and biaxial tests, Figure 2 (Bass et al., 2004; Farshid Ghezelbash et al., 2021).

**Figure 2:**
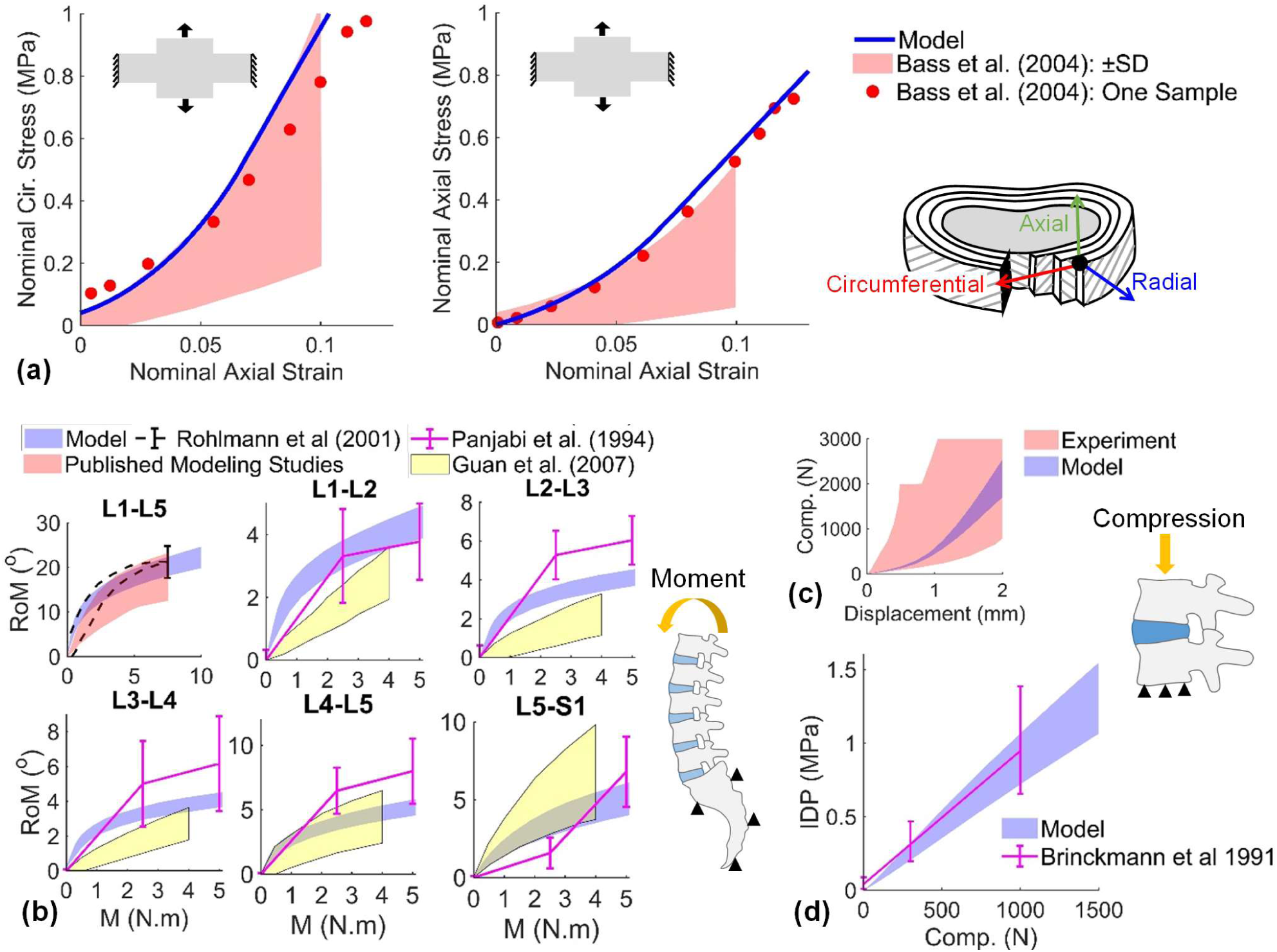
Validation of the subject-specific passive ligamentous spine FE model. (a) Circumferential (Cir.) and axial stress-strain responses of the outer annulus fibrosus tissue model under biaxial loading versus measurements under constrained circumferential strain (circumferential strain=0) (Bass et al., 2004; Hollingsworth and Wagner, 2011); for the specimen size, boundary conditions and stress/strain evaluation, see (Farshid Ghezelbash et al., 2021). (b) Sagittal range of motion (RoM) under flexion moment versus reported *in vitro* measurements of the L1-L5 or L1-S1 lumbar spine segments (Guan et al., 2007; Panjabi et al., 1994; Rohlmann et al., 2001) and other model predictions (Dreischarf et al., 2014). (c) Compression-displacement response of the L2-L3 disc-body-unit model against experimental studies (Ghezelbash et al., 2020a). (d) Estimated intradiscal pressure (IDP) at the L2-L3 disc model versus single-level measurements in compression (Brinckmann and Grootenboer, 1991). To compare to reported cadaver studies, personalized FE-MS model inputs were varied here in the range of body height = 160 – 190 cm, BMI = 19 – 30 kg/m^2^, age = 45 years, and sex = female and male.

#### Passive spine

To evaluate the performance of the passive FE model and the scaling algorithm, displacement response of the L1-S1 ligamentous spine under flexion moment and of the L2-L3 disc body unit in compression as well as the L2-L3 intradiscal pressure-compression were assessed against available *in vitro* experimental studies for different sex, height and weight inputs (Brinckmann and Grootenboer, 1991; Guan et al., 2007; Panjabi et al., 1994; Rohlmann et al., 2001; Anderson et al., 2012b; Ghezelbash et al., 2020a).

#### Intradiscal pressure

Estimated L4-L5 intradiscal pressures were compared with available *in vivo* measurements during various physical activities by considering participants’ height, weight, age and sex (Sato et al., 1999; Takahashi et al., 2006; Wilke et al., 2001).

#### Muscle activity

Estimated muscle activities of an average subject (with mean of kinematics and anthropometric parameters of 19 subjects) during forward flexion without and with a 6 kg hand-load were compared with recorded surface electromyography measurements (Ghezelbash et al., 2020b).

### 2.3 Applications

The subject-specific model was applied to evaluate the effects on spine biomechanics of wearing an exoskeleton and two surgical interventions.

#### Exoskeleton

An exoskeleton system was simulated by three shear deformable elastic beams (each with 30 3-node elements; diameter = 4 mm^2^; elastic modulus = 5 GPa; matching mechanical properties of the system) which were coupled with vertebral rotations at their inserted levels S1 and T12 (Koopman et al., 2020; Näf et al., 2018). The FE-MS model was initially scaled in accordance with the average participants’ anthropometric parameters (Koopman et al., 2020), and subsequently driven by measured mean spine kinematics with/without exoskeleton during forward flexion with a 10 kg hand-load (Koopman et al., 2020; Näf et al., 2018).

#### Surgical interventions

To demonstrate the versatility of the proposed model, we simulated a total nucleotomy intervention at the L5-S1 disc, as well as a spinal fusion procedure (i.e., interbody cage fusion with bi-lateral posterior instrumentation) at the L4-L5 segment assuming a rigid interface connection to fully stabilize the segment (Ebrahimkhani et al., 2021). For fusion simulations, spine kinematics were applied based on *in vivo* imaging studies (Nie et al., 2019). In an additional case, foregoing estimated loads (i.e., shear, compression, and moment) were applied to a more detailed sub model of the stabilized L4-L5 segment to further investigate the effects of the spinal fusion system. Details about the modeling approach are presented in the Supplementary Materials.

## 3 Results

In validation cases, tissue level stress-strain responses of the outer annulus fibrosis computed under biaxial tests had an overall satisfactory agreement with reported measurements, Figure 2a. Global (L1-L5) and segmental (L1-L2 to L5-S1) ranges of motion of the ligamentous FE model under pure flexion moment were found within those of in vitro measurements (Figure 2b). Likewise, compression-displacement (Figure 2c) and compression-intradiscal pressure (Figure 2d) estimations of motion segments had satisfactory agreements with reported measurements. In the passive spine FE model, changes in the model inputs (i.e., sex, height and weight) yielded variations in output responses that were found comparable to those in experiments (Figure 2). For the MS model validation, we compared estimated L4-L5 intradiscal pressures as well as muscle activities against corresponding in vivo measurements collected during various sagittally symmetric tasks, and in both cases, predictions agreed well with recorded experimental results (Figure 3).

**Figure 3:**
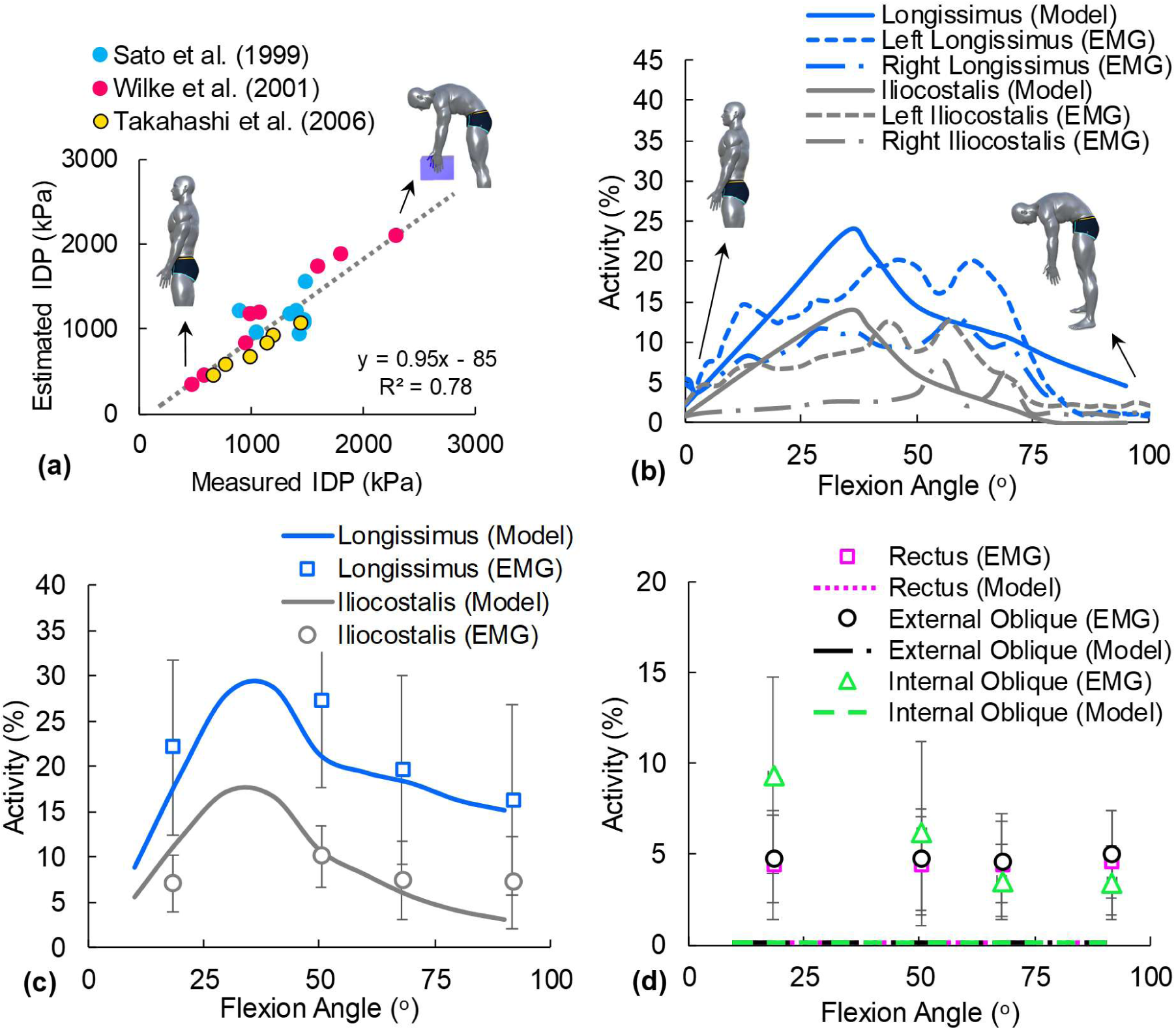
Validation of the subject-specific model of trunk. (a) Estimated intradiscal pressures (IDP), as illustrated by the dotted line, against *in vivo* measurements during various sagittally-symmetric tasks. (b) Comparison between estimated activities of the right and left longissimus pars thoracic and iliocostalis pars thoracic muscles with normalized measured EMG signals during forward flexion of a single subject with no load (body height = 169 cm; body weight = 67 kg; age = 27 years) (Ghezelbash 2016). (c and d) Estimated muscle activities (of a subject with averaged anthropometric parameters and kinematics) against measured EMGs of 19 subjects (error bars represent standard deviation) during forward flexion with a 10 kg hand-load at different flexion angles (i.e., load heights: upright, mid-femur, knee and mid-tibia levels) (Ghezelbash et al., 2020b). R: Rectus; EO: External Oblique; IO: Internal Oblique; M: Multifidus; I; Illiocostalis; L: Longissimus.

In application cases, wearing an exoskeleton (as an ergonomic application) noticeably reduced intradiscal pressure (e.g., 2.2 MPa to 1.9 MPa at the L5-S1 disc) and the von Mises stress in the annulus fibrosus (e.g., 2.9 MPa to 2.1 MPa at the L4-L5 disc). Furthermore, contact areas and stresses at facet joints decreased, particularly at the upper T12 and L1 levels, when wearing the exoskeleton (Figure 4). Exoskeleton beams resisted in total 6.3 N.m moment (20 N shear forces), which led to smaller forces in global (746 N versus 774 N) and local (2011 N versus 2236 N) muscles at 75° forward flexion.

**Figure 4:**
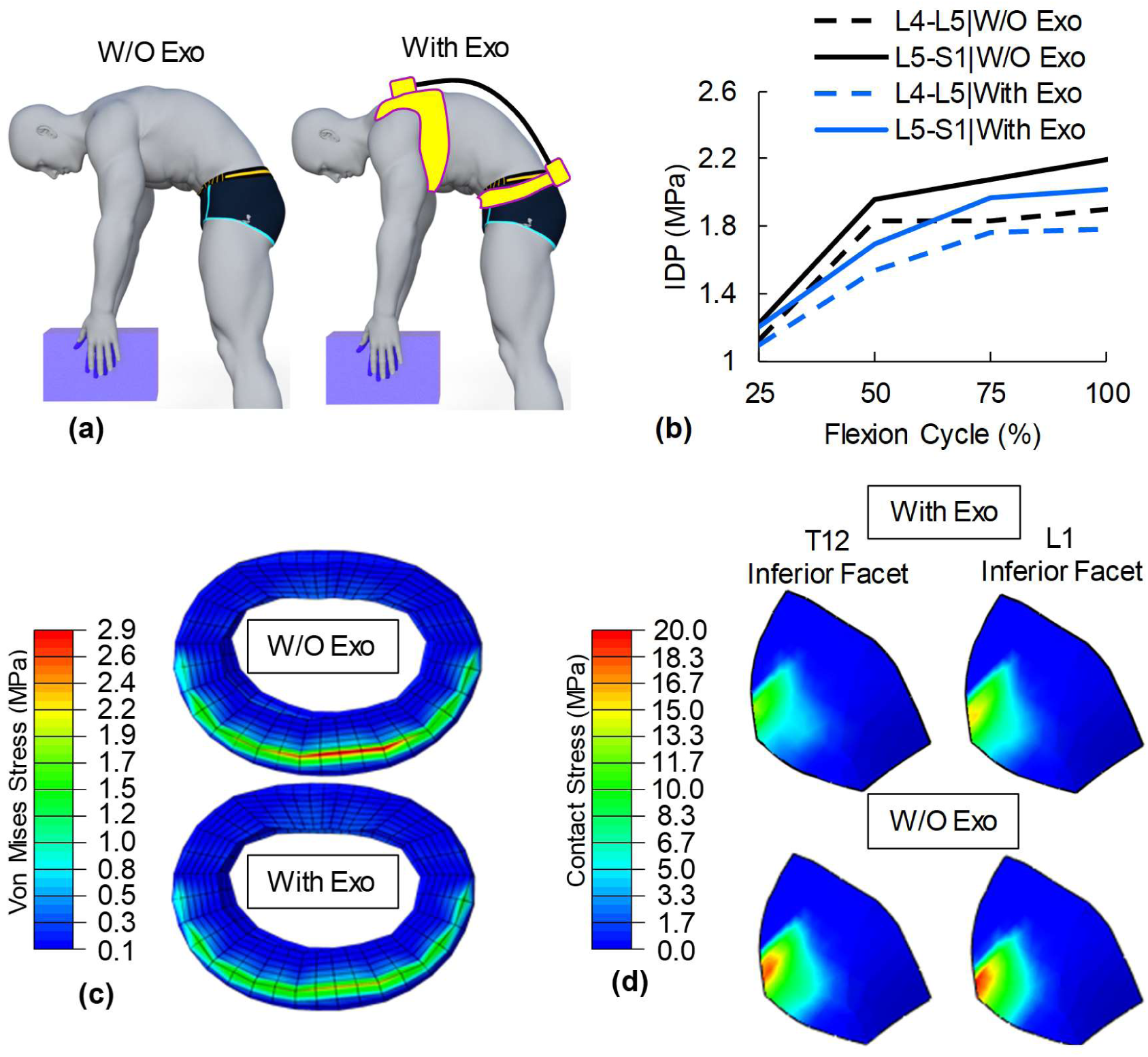
The effects of (a) wearing an exoskeleton (Exo) on (b) intradiscal pressure (IDP) at two lowermost disc levels during forward flexion, (c) annulus fibrosus matrix von Mises stress (L4-L5 disc) at full flexion, and (c) articular facet contact stresses (at full flexion; right inferior facets) under a 10 kg hand-load. Model inputs were chosen based on the reported mean anthropometric parameters of subjects (Koopman et al., 2020).

In clinical applications, spinal fusion increased the intradiscal pressure in both adjacent discs (L3-L4: 1.41 MPa versus 1.17 MPa; L5-S1: 1.72 MPa versus 1.58 MPa at 50° flexion with a 10 kg hand-load), but nucleotomy had a minimal effect on the estimated intradiscal pressures elsewhere away from the L5-S1 disc (Figure 5). Following nucleotomy, the inner annulus layers at the denucleated L5-S1 disc bulged inward exhibiting larger matrix radial strains and von Mises stresses (6 MPa versus 3.8 MPa). In all cases (i.e., normal, nucleotomy and spinal fusion), the maximum collagen fiber strain at the L5-S1 disc reached the peak of 13%; however, in the spinal fusion case, a larger portion of disc collagen fibers experienced such large strains. Simulated single-level model of the stabilized L4-L5 segment indicated that the posterior rods as well as vertebrae experienced rather large stresses (peaks of 380 MPa and 129 MPa, respectively).

**Figure 5:**
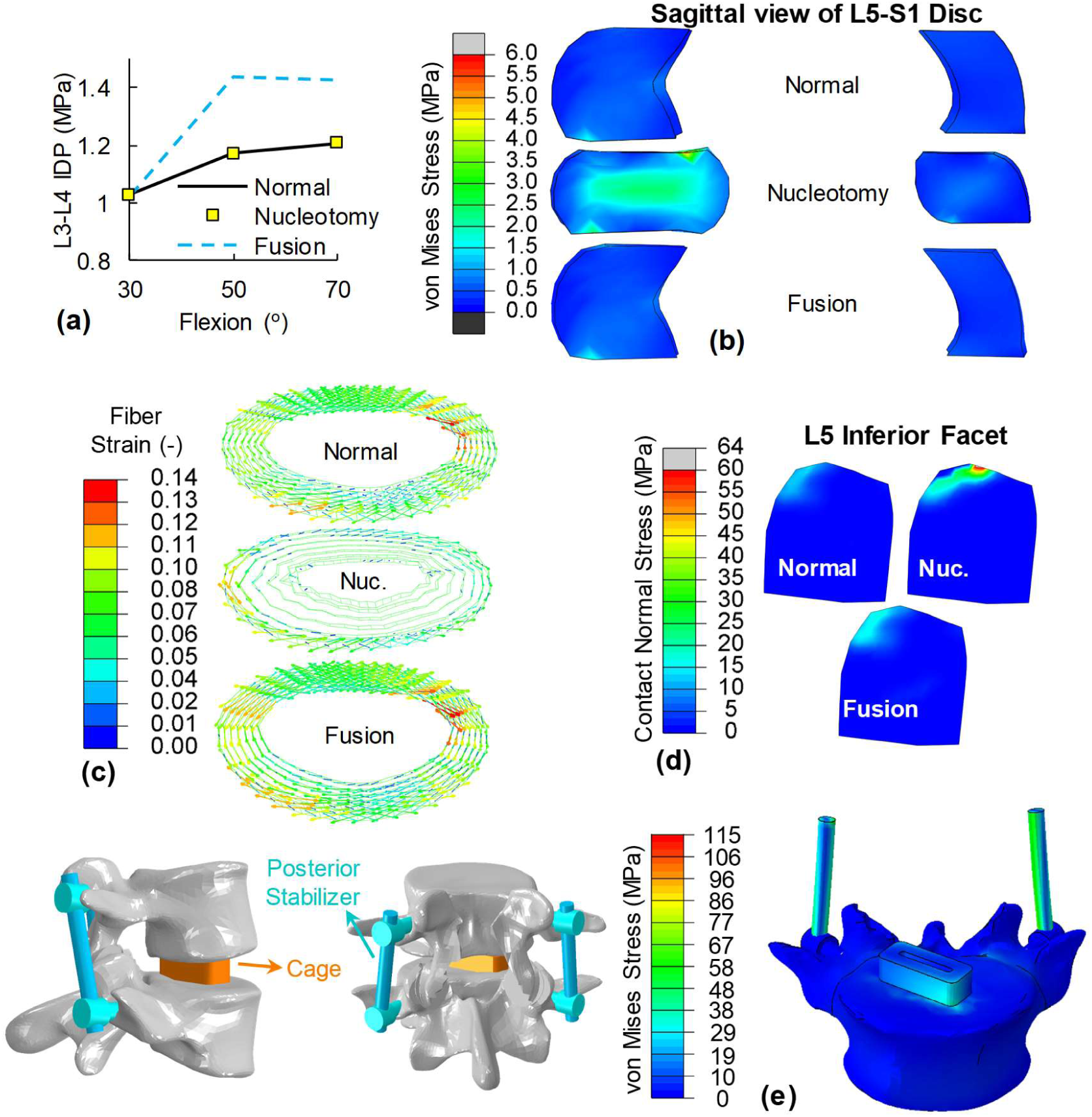
The effects of different surgical interventions (total L5-S1 nucleotomy – Nuc. – and L4-L5 fusion) on spine biomechanics: Estimated (a) L3-L4 intradiscal (IDP) pressure, (b) von Mises stress at the L5-S1 disc annulus in the sagittal plane, (c) fiber strains at the L5-S1 disc, and (d) contact stress at the L5 inferior right facet joint during forward flexion with 10 kg hand load, (e) predicted von Mises stress in the L4-L5 fusion single-level model (external loads were applied from the integrated subject-specific model). Unless indicated otherwise, presented parameters are for 70° forward flexion. Model inputs were set at body height: 173 cm, body weight: 73 kg, age: 45 years, and sex: male.

## 4 Discussion

We developed and validated a subject-specific coupled FE-MS model that incorporates both passive and active trunk components in a single model. The scaling algorithm individualized the detailed ligamentous spine (e.g., disc height/area, lordosis, ligament insertions, facet locations), gravity loads/locations as well as the muscle architecture based on the subject’s height, weight, age and sex. The model had a satisfactory performance in predicting responses at the tissue-level, disc-level, entire lumbar spine as well as the MS trunk. To showcase the capabilities of the individualized FE-MS model in ergonomics and clinical cases, we explored the effects of wearing an exoskeleton and two surgical interventions on spine biomechanics.

### 4.1 Limitations

Spinal tissues and mechanical responses were assumed to be elastic and time-independent (Khalaf and Nikkhoo, 2021; Zehr et al., 2020; Zheng et al., 2022); nevertheless, we simulated intervertebral disc as an incompressible elastic material which could accurately represent the short-term response of a poroelastic model (Ateshian et al., 2006; Ghezelbash et al., 2022; Shirazi et al., 2008). The scaling algorithm took account of geometric changes in the passive spine as well as the muscle architecture; however, we did not adjust changes in the intrinsic material properties with model inputs (as expected in general population and also with ageing and disc degeneration) because of the complexity of the issue as well as the paucity of the data in the literature. In all simulations, therefore, we intentionally avoided the consideration of individuals older than 45 years. An optimization algorithm was used to estimate muscle forces; thus, coactivity in antagonist abdominal muscles were not considered (Kian et al., 2019). Due to the lack of data and geometric complexity involved, the scaling algorithm did not modify geometrical features of facet joints (i.e., facet articular shapes and gaps) in the reference model which was originally reconstructed from computed tomography images of a cadaver lumbar spine (Breau et al., 1991); however, it should be noted that some facet parameters (such as facet gap) can potentially affect spine biomechanical response (Bashkuev et al., 2020; Schmidt et al., 2013; Shirazi-Adl, 1994b). It should be noted that this framework could be combined with automated imaging segmentation and resolve foregoing issues (Matos et al., 2023; Pinheiro et al., 2018; Zhang and Wang, 2021). The layerwise approach (i.e., distinctively simulating each lamella) is a more anatomically accurate method of simulating the annulus fibrosus; nevertheless, we used the homogenized approach (i.e., combing adjacent layers into a single layer) both to reduce the computational burden and to concur with our earlier study showing that both approaches estimated similar global responses (e.g., compression-displacement) (Farshid Ghezelbash et al., 2021). Also for the sake of computational efficiency, each of T12-L5 vertebral bodies was simulated as a collection of an anterior and a posterior rigid body interconnected by two deformable beams at their respective pedicles to represent the vertebral deformities expected under larger loads. If stresses in vertebral bodies are, however, of interest, then an isolated model of a single or multi-motion segments with vertebrae represented as deformable bodies would be required as considered here in the fusion simulation model (Figure 5e). Finally, while changes reported in lumbar segmental lordosis angles and hence in the entire lumbar lordosis were locally simulated (Tang et al., 2022), the global effects on the position of T1 and the rest of spine such as facet orientations-gaps were overlooked.

### 4.2 Validation

In this study, we validated responses of our personalized FE-MS model by comparison with available in-vitro and in-vivo measurements at both tissue and global levels of the model (e.g., annulus fibrosus under biaxial loading, ranges of motion under flexion moment, compression-displacement of discs, IDP and EMG). Overall, all computed responses were in a satisfactory agreement with available measurements (Figures 2 and 3). It should be noted that a large scatter in both predictions and measurements exists due to inherent inter-specimen and inter-individual variabilities as well as due to limitations in and parameter-dependence of model predictions.

### 4.3 Novelties

Unlike conventional MS models which use simplistic components (such as spherical joints with/without linear/nonlinear rotational springs or beams) to take account of the passive ligamentous spine; here, we have used a detailed FE model of the spine which was individualized through a novel scaling algorithm. Such an integrated approach provides us also with much-needed outputs such as the strain/stress fields in discs/facets/ligaments which are essential for a comprehensive risk analysis and a remarkable flexibility to realistically simulate various surgical procedures. For instance, conventional MS models cannot simulate a nucleotomy procedure or evaluate the risk of a ligament tear or annulus fibrosus failure simply because they do not distinctly simulate intervertebral discs, facet joints and ligaments. Isolated passive FE models, on the other hand, fundamentally fail to mimic realistic loading conditions by overlooking muscles as active components. Passive FE spine models often use idealized loading (such as a pure moment with/without a follower load) to account for complex physiological loading conditions. Additionally, we developed a novel scaling algorithm to adjust various geometric features of the model, and our analyses demonstrated that changes in model inputs (i.e., body height, body weight and sex) could replicate the variations in experimental data (Figure 2b, c and d), confirming that a part of the scatter in recorded responses is explained by morphologic changes.

Finally, the current integrated model accurately accounts for the continuous changes in the passive stiffness properties under varying external loads (Shirazi-Adl, 2006) that are due primarily to the activation in muscles. Alterations in the passive properties, on the other hand, influence the required activation level in muscles. Such realistic simulation of passive-active coupling is possible only when both components are accurately represented alongside in a model study. The coupled model of the lower extremity for the knee (Makani et al., 2022; Sharifi and Shirazi-Adl, 2021) and hip (Hua et al., 2022) attest to this point.

### 4.4 Applications

The current model can be employed to carry out a comprehensive neuromuscular risk analysis, surgical intervention planning as well as the evaluation of assistive devices such as exoskeletons. We investigated the effects of wearing an exoskeleton on spine biomechanics and predicted that it can reduce compression loads on spine (by 15%), matrix stresses in annulus fibrosus (by 27%), facet joint contact stresses (by 20%) and intradiscal pressures (by 14%), which altogether can potentially decrease the risk of spinal injuries. It is to be noted that a conventional model can only provide insights on muscle activities along with global spine shear/compression loads. Finally, although we found that using an exoskeleton unloads the spine, future studies should explore likely alterations in the load-transfer pathways in presence of such assistive devices; particularly in the lower extremity hip and knee joints.

We simulated nucleotomy as well as spinal fusion interventions using the individualized integrated model. Total nucleotomy procedure did not alter intradiscal pressures in the remaining intact levels but markedly affected segmental load-sharing (Figure 5) at the denucleated level itself. Nucleotomy caused a substantial inward disc bulge, a marked increase in annulus radial strain and a much larger matrix stress as well as a load transfer from the disc to facets (max stress: 62 MPa versus 33 MPa). These findings suggest both a higher risk of injury and overload to facet joints and an annulus inward instability and circumferential tears and hence disc degeneration as the disc nucleus loses its fluid content and internal pressure.

On the other hand, spinal fusion increased intradiscal pressure in the adjacent levels (L3-L4: 1.41 MPa versus 1.17 MPa; L5-S1: 1.72 MPa versus 1.58 MPa). This procedure perturbs the activation level in muscles as well as the relative contribution of passive tissues at various levels. Under the same overall flexion angle and reported larger pelvic rotation (Nie et al., 2019), smaller lumbar rotation increases the contribution of muscles (as active components) to counterbalance external moments, which leads to greater intradiscal pressures and larger collagen fiber strains (Figure 5). These effects are however dependent on the prescribed rotations following fusion surgery.

## 5 Conclusions

In this study, we developed a modeling framework that overcomes the limitations of previous fully passive and conventional MS models by combining a detailed passive spine representation with a complex active muscle architecture, resulting in a parametric model that is scalable based on individual factors. This model has been rigorously validated at multiple levels: tissue-level responses, each functional level of the spine, and the integrated model (i.e., muscle activities, intradiscal pressures), thereby confirming its validity. Distinct from MS models (which are incapable of simulating surgical interventions), and from passive spine models (that do not incorporate realistic muscle loads), our framework has demonstrated its versatility by simulating a range of surgical interventions such as nucleotomy and fusion. Unlike current approaches, this novel framework can provide detailed tissue-level responses, which are critical for a comprehensive injury analysis, and we have shown its performance in evaluating the effect of wearing an exoskeleton. This approach holds promise for enhancing personalized medicine by aiding in surgical planning, and it can contribute to the design of preventative measures in the workplace, ultimately aiming to reduce the risk and incidence of spine-related injuries.

## 6 Acknowledgement

This study was supported by the Institut de recherche Robert-Sauvé en santé et en sécurité du travail (IRSST; 2019-0018), Mitacs Elevate Fellowship (Mitacs-IT18290), Fonds de Recherche du Québec Nature et technologies (FRQNT), and Natural Sciences and Engineering Research Council of Canada (CRSNG RGPIN-03356).

## 8 Supplementary Materials

### 8.1 L4-L5 Sub-Model

In the musculoskeletal model, all vertebrae were simulated as a collection of two rigid bodies attached by two flexible beams at their pedicles. To evaluate the stress distribution at the L4-L5 level, a finite element sub-model was employed. Predicted resultant moments and forces from the detailed musculoskeletal model were applied at the L4 vertebra (through a rigid plate attached to the upper endplate at L4), and the L5 vertebra was fixed (at its lower endplate plate). In the fusion simulation, the nucleus pulpous as well as a part of the annulus fibrosus (posterolateral region) were removed. A polyetheretherketone (PEEK) cage was diagonally placed within the available disc space and filled with bone graft. A contact interaction between the cage and adjacent bones with a friction coefficient of 0.2 was assumed. A spinal stabilizer system (pedicle screws, diameter of 6.5 mm, and rods, diameter of 5.5 mm) was assembled in the model, and screws were inserted through the pedicle and embedded in the vertebral body assuming attached interfaces.

**Figure S1:**
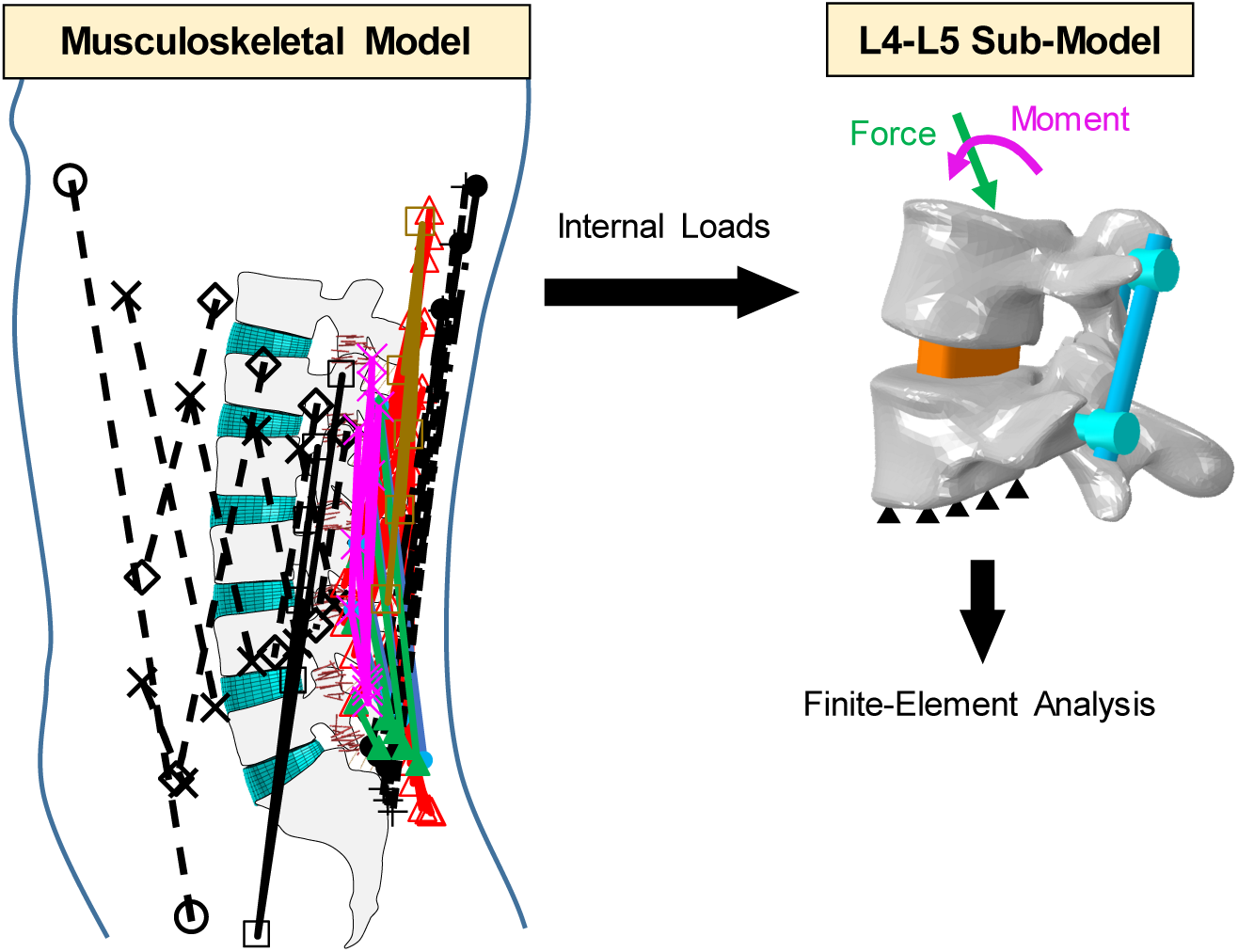
Internal loads on the isolated L4-L5 sub-model were estimated from the musculoskeletal model.

**Table S1:** Material properties of the finite element-musculoskeletal model as well as the L4-L5 sub-model.

| Component | Material property | Reference |
| --- | --- | --- |
| Annulus fibrosus matrix | Incompressible Mooney-Rivlin hyperelastic model ($C_1 = 0.14$ MPa; $C_2 = 0.035$ MPa) | (F Ghezelbash et al., 2021) |
| Annulus fibrosus collagen fiber | Nonlinear stress-strain curves | (Shirazi-Adl et al., 1986) |
| Ligaments | Nonlinear stress-strain curves | (Shirazi-Adl, 1994c; Shirazi-Adl et al., 1986) |
| Cortical bone | Linear elastic ($E = 12000$ MPa; $\nu = 0.3$) | (Goto et al., 2003; Schmidt et al., 2007b) |
| Cancellous bone | Linear elastic ($E = 200$ MPa; $\nu = 0.315$) | (Goto et al., 2003; Schmidt et al., 2007b) |
| Posterior elements | Linear elastic ($E = 3500$ MPa; $\nu = 0.25$) | (Huang et al., 2016; Shirazi-Adl, 1994c; Shirazi-Adl et al., 1986) |
| Cage | Linear elastic ($E = 3500$ MPa; $\nu = 0.3$) | (Huang et al., 2016) |
| Graft | Linear elastic ($E = 50$ MPa; $\nu = 0.2$) | (Huang et al., 2016) |
| Instrumentation | Linear elastic, titanium ($E = 110000$ MPa; $\nu = 0.2$) | (Huang et al., 2016) |

